# Development Optimization and Validation of RT-LAMP based COVID-19 Facility in Pakistan

**DOI:** 10.1101/2020.05.29.124123

**Authors:** Farhan Haq, Salmaan Sharif, Adnan Khurshid, Imran Shabbir, Muhammad Salman, Nazish Badar, Aamir Ikram, Abdul Ahad, Muhammad Faraz Arshad Malik

**Affiliations:** Department of Biosciences, COMSATS University Islamabad; National Institute of Health, Islamabad; Department of Biotechnology and Bioinformatics, International Islamic University, Islamabad

## Abstract

The pandemic SARS-CoV-2 (Severe acute respiratory syndrome coronavirus 2) has created a widespread panic across the globe especially in the developing countries like Pakistan. The lack of resources and technical staff are causing havoc challenges in the detection and prevention of this global outbreak. Therefore, a less expensive and massive screening of suspected individuals for COVID-19 is required. In this study, a user-friendly technique of reverse transcription-loop mediated isothermal amplification (RT-LAMP) was designed and validated to suggest a potential RT-qPCR alternate for rapid testing of COVID-19 suspected individuals. A total of 12 COVID-19 negative and 72 COVID-19 suspected individuals were analyzed. Both RT-qPCR and RT-LAMP assays were performed for all the individuals using open reading frame (ORF 1ab), nucleoprotein (N) and Spike (S) genes. All 12 specimens which were negative using RT-qPCR were also found negative using RT-LAMP assay. Overall 62 out of 72 positive samples (detected using RT-qPCR) were found COVID-19 positive using RT-LAMP assay. Interestingly all samples (45) having Ct values less than 30 showed 100% sensitivity. However, samples with weaker Ct values (i.e., => 35) showed 54% concordance, suggesting potential false negatives or false positives in RT-LAMP or RT-qPCR results, respectively.

Overall comparative assessment showed that RT-LAMP assay showed strong sensitivity and specificity and can be used as an alternative strategy for rapid COVID-19 testing. Hence, based on fast processing time, minimal risk of specimens transfer and utilizing available resources, LAMP based detection of COVID-19 is strongly advocated especially for developing countries.

## Introduction

Global outbreak of SARS-CoV-2 (Severe acute respiratory syndrome coronavirus 2) is creating a havoc challenge especially in developing countries like Pakistan^1,2^. Being 5^th^ populated country, sharing borders with two adversely affected countries (Iran and China), a massive screening of suspected individuals for COVID-19 is required^3^. Based on variable clinical symptoms ranging from mild flu to life threatening shortage of breath signs it is pertinent to establish an efficient diagnostics tool for earliest detection SARS-CoV-2. So far all commercially COVID-19 tests are relying on two strategies. In the first strategy, variety of immunological assays are developed responsible for detection of antibodies produced by individuals as a result of current or past exposure to the virus^4^. The second strategy focusses on SARS-CoV-2 viral RNA detection using polymerase chain reaction (PCR) ^5^. Additionally, loop mediated isothermal amplification-based techniques are in process ^6^. All these strategies have preferential advantages depending upon the objectives of the screening. As immune assays help to identify individuals retaining developed antibodies against virus and could be potential convalescent plasma donors. Genome based testing facilities help in diagnosis of SARS-CoV-2-infected individuals.

As per National Command and Operation Control (https://ncoc.gov.pk/), there are 94 laboratories offering detection of COVID-19 in Pakistan. All these labs are using real time PCR techniques (RT-qPCR) for detection of SAR-Cov-19. Despite of technique’s robustness, challenges related to increasing number of affected cases in both urban and rural areas, direly require fast and less expensive alternate techniques. Since installation of expensive instrument, prolong reaction time (~3hr), cost of reagents and trained staff availability in an emerging outbreak (COVID-19) are debilitating its effective utilization across Pakistan.

Here, an alternate user-friendly technique of reverse transcription-loop mediated isothermal amplification (RT-LAMP) was designed and validated against COVID-19. Earlier, LAMP assay was used for detection of Hepatitis B virus (HBV)^7^. Briefly, *Bacillus stearothermophilus* (*Bst* polymerase) along with set of 4 primers (2 inner and 2 outer primers) were used in the assay. Reaction began with the binding of the inner primers containing sequences of sense and anti-sense strands of DNA. Here, Bst DNA polymerase retain 5’→3’ polymerase activity while lacking 5’→3’ exonuclease activity for 1 hr at 65°C. Addition of reverse transcriptase along with Bst polymerase in LAMP assay can also enhance its utilization for rapid detection of RNA based viruses. In light of these modifications, LAMP technology was efficiently been advocated by World Health Organisation (WHO) under ASSURED (Affordable, Sensitive, Specific, Userfriendly, Rapid and Robust, Equipment free, Deliverable to end users) criteria ^8^.

## Methodology

### Cohort Identification

National Institute of Health (NIH) is a core national referral resource Institute responsible for accreditation and testing of COVID-19 screening facilities. For the present study, 72 COVID-19 suspected individuals were enrolled at NIH. These individuals were categorized into different groups based on the Ct values retrieved from RT-qPCR.

### Specimen Collection and RNA Isolation

Nasopharyngeal specimens from these individuals were collected at designated specimen collection facility. After immersing these specimens in virus transfer buffer, these tubes were processed further for RNA isolation using commercially available kit. TANBead Nucleic Acid Kit was used for viral RNA isolation by using TANBead Nucleic Acid Extractor (model SLA-16/32) as per manufacturer instructions. Briefly, this kit used silicon dioxide layer coated magnetic beads for isolation of nucleic acid in 40 min.

### Designing of RT-LAMP assay Primer

Primers targeting open reading frame (ORF 1ab), nucleoprotein (N) and Spike (S) genes were designed using Primer Explorer V5 software. As per guidelines of LAMP assay, these panel of primers were specifically designed against a predefined genomic region of 2019-nCoV (genome Reference Sequence: NC_045512.2). Primer sequences will be provided on request.

### Reaction Conditions for RT-LAMP Assay

WarmStart Colorimetric LAMP 2× Master Mix (New England Biolabs) vials were used for RT-LAMP assay. Three separate primer mix for each gene were prepared as working stock vials for the reaction. Reaction tube for each specimen containing 2× master mix (12.5 μl), 10× primer mix (1×; 2.5 μl), RNA (2 μl) and water (8 μl) were left at thermomixer® (Eppendorf) for incubation. Efforts were made to set forth best optimum temperature ranging from 62C° −65C° with time ranging from 45 min-75min for better yield.

### Estimation of assay Sensitivity and Specificity

Being a colorimetric LAMP assay, change in color of reaction mix from pink to yellow color was set as indicator of presence of virus. Since generation of concatemers result in lowering of pH from basic to acidic state. Hence, both amplification and detection of respective concatemers were completed in *a single step*. In addition, samples were also validated using gel electrophoresis results.

### QPCR based Detection of COVID-19 for Respective Specimens

All aforementioned specimens (72) were analyzed using available COVID-19 diagnostic kits. RNA sample of these respective specimens were processed for diagnosis using RT-qPCR kit as per manufacturer instructions. Hence RT-LAMP findings were also cross checked with available gold standard testing facility of RT-qPCR.

## Results

### Identification of COVID-19 in the cohort

Majority of these individuals were male expatriate Pakistani returning home with median age of 38Yrs. All 12 specimens which were negative using RT-qPCR were also found negative using RT-LAMP assay.

### Detection of RT LAMP based COVID-19

Overall 62 out of 72 positive samples (detected using RT-qPCR) were found COVID-19 positive using RT-LAMP assay. After extensive optimization, RT-LAMP reaction optimum conditions for all three genes were set at 65C° for 1hr duration. COVID-19 positive samples showed pronounced change in coloration from pink to yellow color against all three respective genes (Figure 1).

**Figure 1:**
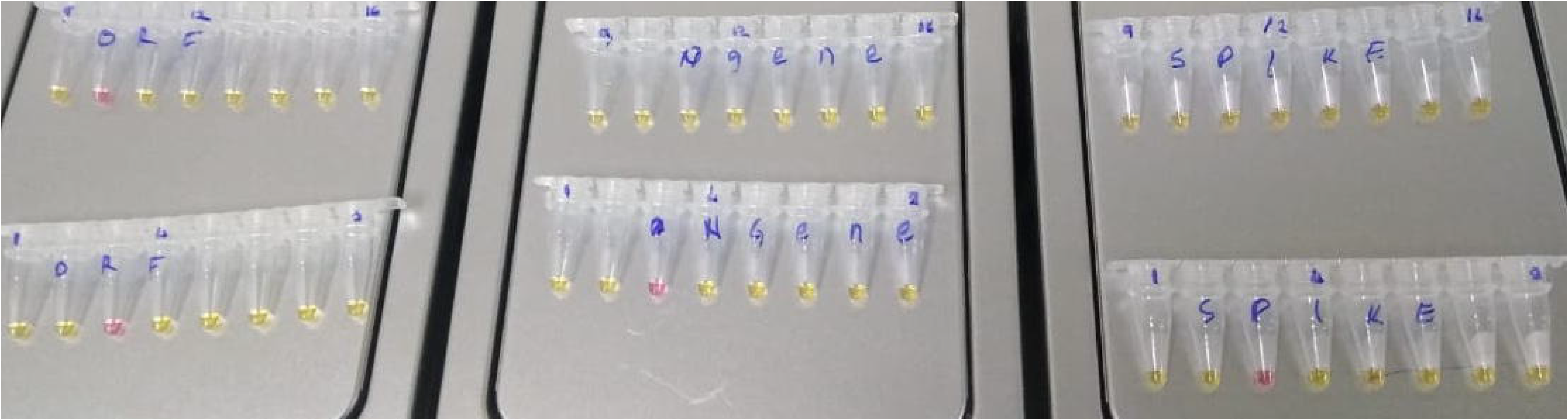
Representative image of colorimetric RT-LAMP assay results of COVID-19 patients for open reading frame (ORF 1ab), nucleoprotein (N) and Spike (S) genes, respectively. Yellow is the indicator of presence of virus.

### Comparative assessment of qPCR and RT-LAMP based COVID-19 detection

Next, we divided positive samples into different RT-qPCR groups based on their Ct values. Interestingly all samples (45) having Ct values less than 30 showed 100% sensitivity (Figure 2). 10 out 14 samples with Ct values < 35 showed positivity using RT-LAMP assay. However, 7 out of 13 samples with Ct values => 35 showed positivity using RT-LAMP assay. This suggests that samples with Ct > 35 should be re-evaluated for potential false positivity of RT-RT-qPCR (Figure 2).

**Figure 2:**
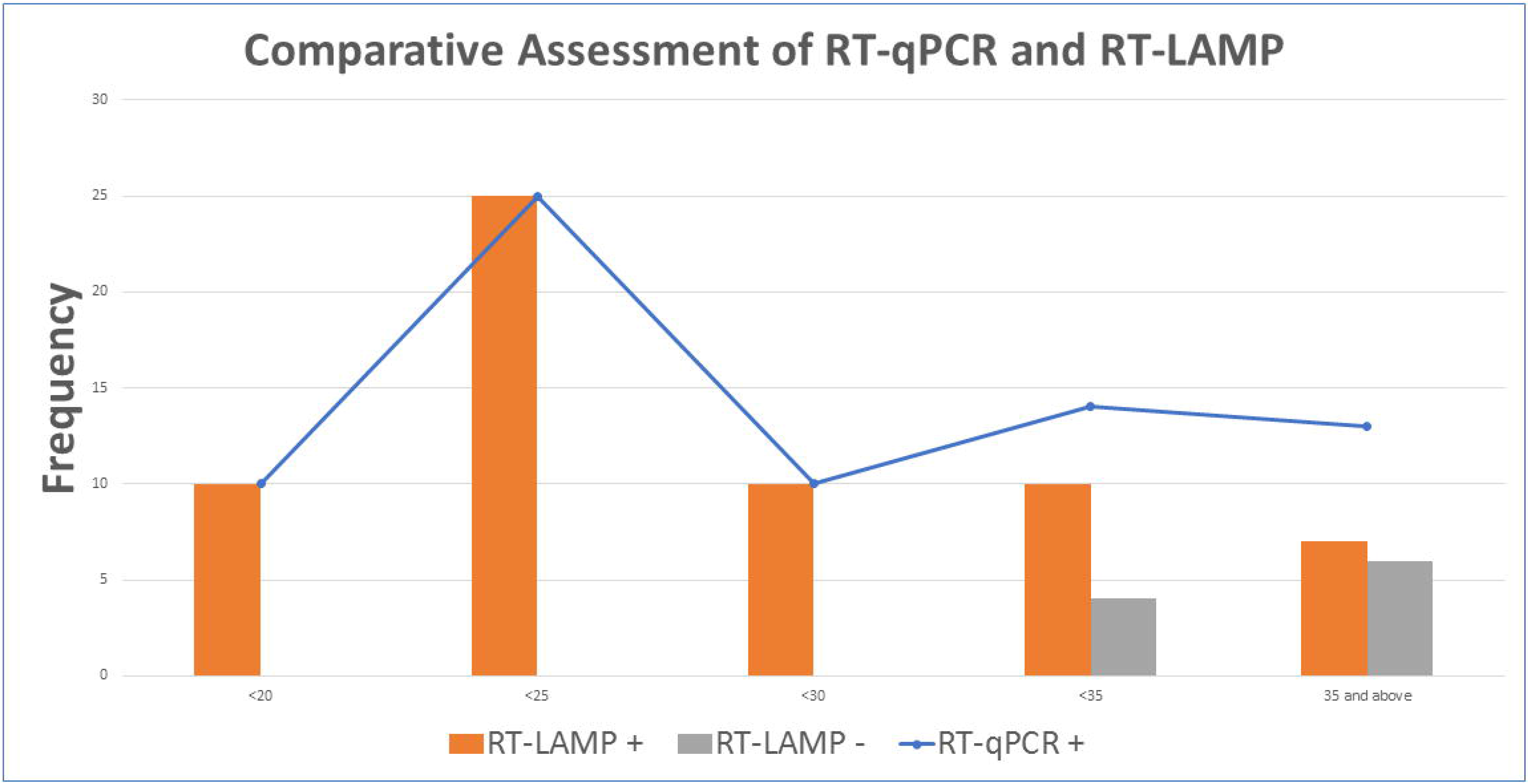
Comparative assessment of RT-qPCR and RT-LAMP based COVID-19 detection.

## Discussions

Overall comparative assessment showed that RT-LAMP assay showed strong sensitivity and specificity and can be used as alternate strategy for rapid COVID-19 testing. Earlier, LAMP assay been widely used for screening of food pathogens ^9^, detection of avian influenza virus ^10^ and gastric cancer metastasis ^11^. Interestingly, several modifications towards earliest identification of concatemers also improve and largely reduced assay dependency on expensive instruments. Conventionally, LAMP products could also be detected by using agarose gel electrophoreses after 1hr incubation at high temperature ^7^. Later on, formation of white precipitate (magnesium pyrophosphate) in the reaction tube may also traced either by naked eye or spectrophotometry^12^. Pyrophosphate binding with a divalent metallic ion magnesium ion (Mg++) was the principle underlying the development of turbidity where increase turbidity was directly correlated with DNA synthesized. Alternatively, introduction of calcein (fluorexan) also acting as fluorescent metal indicator was used to enhance visual detection of LAMP reaction.

In the beginning of reaction, binding affinity of manganous ion (Mn2+) was responsible for quenching of calcein signals. During amplification of targeted reaction, generated pyrophosphate forming complex with Mn2+, facilitate calcein release resulting in fluorescence detected under UV ^13^. Similarly, ethidium bromide and Malachite Green dyes have also been used as detection for the LAMP amplification products ^14^. In Warmstart mastermix, pH based approach was used where release of pyrophosphate result in lowering of pH from pink to yellow. This colorimetric assay is potentially the easiest approach where we can have reduced assay dependency on expensive instruments as well as highly trained technicians.

Currently, few LAMP based kits like ID NOW COVID-19 and iAMP COVID-19 also approved by FDA are commercially available for SARS-Cov-2 detection. ID NOW COVID-19 (Abbott Diagnostics) target isothermal amplification of a single gene (RdRP) with fast processing time (~13min). However, it can process one sample per run with close dependency on reagents and consumables as per manufacturer instructions. iAMP COVID-19 (Atila BioSystems) target isothermal amplification of ORF1ab and N genes with processing time of 1.5hrs. Based on global disease burden, availability of these kits at Pakistan would be extremely challenging especially when export of certain kits are barred from country of origin (like Loopamp 2019-nCOV Detection Kit of Eiken export from Japan). In the current study, detection of SARS-Cov-2 utilizing 3 genes (ORF1ab, N and S) genes immensely improved LAMP test detection threshold and reliability on inhouse resources. Hence, based on fast processing time, minimal risk of specimens transfer and utilizing available resources, LAMP based detection of COVID-19 is strongly advocated for developing countries.

## Acknowledgements

We are grateful to National Institute of Health, COMSATS University Islamabad and Higher Education for providing needful resource facilitation, training and funding resource. All authors declare no conflict of interest.

## Authors Approval

All authors have seen and approved the manuscript, and that it hasn't been accepted or published elsewhere.

## Notes

### Competing Interest Statement

The authors have declared no competing interest.

